# The endoplasmic reticulum membrane protein complex localizes to the mitochondrial - endoplasmic reticulum interface and its subunits modulate phospholipid biosynthesis in *Trypanosoma brucei*

**DOI:** 10.1101/2021.06.17.448810

**Authors:** Advaitha Iyer, Moritz Niemann, Mauro Serricchio, Caroline E. Dewar, Silke Oeljeklaus, Luce Farine, Bettina Warscheid, André Schneider, Peter Bütikofer

## Abstract

The endoplasmic reticulum membrane complex (EMC) is a versatile complex that plays a key role in membrane protein biogenesis in the ER. Deletion of the complex has wide-ranging consequences including ER stress, disturbance in lipid transport and organelle tethering, among others. Here we report the function and organization of the evolutionarily conserved EMC (TbEMC) in the highly diverged eukaryote, *Trypanosoma brucei*. Using (co-) immunoprecipitation experiments in combination with mass spectrometry and whole cell proteomic analyses of parasites after depletion of select TbEMC subunits, we demonstrate that the TbEMC is composed of 9 subunits that are present in a high molecular mass complex localizing to the mitochondrial-endoplasmic reticulum interface. Knocking out or knocking down of single TbEMC subunits led to growth defects of *T. brucei* procyclic forms in culture. Interestingly, we found that depletion of individual TbEMC subunits lead to disruption of *de novo* synthesis of phosphatidylcholine (PC) or phosphatidylethanolamine (PE), the two most abundant phospholipid classes in *T. brucei*. Downregulation of TbEMC1 or TbEMC3 inhibited formation of PC while depletion of TbEMC8 inhibited PE synthesis, pointing to a role of the TbEMC in phospholipid synthesis. In addition, we found that in TbEMC7 knock-out parasites, TbEMC3 is released from the complex, implying that TbEMC7 is essential for the formation or the maintenance of the TbEMC.

## Introduction

The endoplasmic reticulum (ER) is a vast multifarious membrane system that plays a key role in lipid and protein synthesis in a cell. It is known to house many protein complexes that aid its function, including the conserved ER membrane protein complex (EMC)^1^. The EMC was first described more than a decade ago in *Saccharomyces cerevisiae* as a complex composed of 6 proteins^2^ and is now believed to contain 8 subunits localizing exclusively to the ER^3^. Since then, EMCs have been identified in every major eukaryotic lineage^1^. The wide distribution of the EMCs and the high degree of sequence conservation between the corresponding subunits of the complex underline their functional importance. Currently, the primary role of the EMC is in protein insertion into the ER membrane. Its depletion causes pleiotropic effects, including ER stress, viral protein destabilization and alterations in lipid transport^2,4–8^. However, many of these effects may ultimately be due to failed insertion of ER membrane proteins.

Early work in *S. cerevisiae* implied that the EMC tethers the ER to the mitochondria^7^. Simultaneous deletion of multiple subunits of the EMC lead to defects in phospholipid trafficking from the ER to the mitochondria, resulting in impaired mitochondrial function^7^. Subsequently, the role of the EMC in phospholipid metabolism has received little additional attention. Instead, more recent studies have focused on the role of the EMC in the biogenesis of multipass membrane proteins^9^ and the association of its subunits with cell surface membrane proteins^10–12^, indicating that it acts as transmembrane domain (TMD) chaperone^4^ or insertase^13^. In mammalian cells, insertion of the ER resident enzyme, squalene synthase, was found to depend on a functional EMC^13^ and in reconstitution experiments, a purified 10 component EMC facilitated membrane insertion of squalene synthase and other reporter proteins, indicating that the EMC mediates insertion of proteins with low hydrophobicity^13^. Furthermore, a mass spectrometry (MS)-based proteomic analysis identified a list of EMC subunit-dependent proteins containing TMDs with polar and/or charged residues^14^. In structural studies, the human EMC has been suggested to mediate membrane insertion of proteins containing positively charged amino acids within their TMD^15^. The model implicates EMC3 in the capture and targeting of the substrate to the membrane. Subsequently, substrate insertion may be achieved via a hydrophilic vestibule composed of EMC3 and EMC6, whereby a local thinning of the membrane and a positively charged TMD patch in the bilayer further assist substrate insertion. The unstable binding of EMC3 and EMC6 with the substrate allows its dissociation from the EMC. Finally, the substrate encounters the lumenal EMC1, which has been proposed to act as a recruitment site for chaperones and co-factors involved in protein folding^15^.

The EMC’s role in membrane protein insertion may also explain, at least in part, its involvement in viral protein expression and replication by interacting with TMDs of viral proteins that are crucial for viral replication, contributing to their stabilization, folding and expression^8^. In human cells, the absence of the EMC caused a subset of membrane proteins to be inaccurately expressed and integrated into membranes, leading to a substantial reduction in infection rate of the viruses^8^.

Reconstructions of the human EMC revealed a 10 member complex with a tripartite configuration^15^. Based exclusively in the ER, EMC1, EMC4, EMC7 and EMC10 form an L-shaped lumenal region, with EMC2 and EMC8 anchoring a basket shaped cytosolic region and a total of 12 TMDs making up the membrane spanning region in the ER^15^. Additional reports revealed that EMC1, EMC2, EMC3, EMC5 and EMC6 form the core while EMC4, EMC7 and EMC10 constitute peripheral components of the complex^4,16^.

Analyses by homology searching algorithms have deduced that the EMC was present in the last eukaryote common ancestor (LECA)^1^, making the study of EMCs an interesting pursuit apropos the evolution of ancestral ER proteins. Experimental studies on the EMC have so far been limited to the Opisthokonts ^17^, except for one report in *Trypanosoma brucei*, which belong to the eukaryotic supergroup of Excavata. The study showed that depletion of EMC2 in *T. brucei* bloodstream forms lead to increased cell doubling time and resistance to isometamidium, berenil, and acriflavine-induced kinetoplast loss^18^. However, the molecular organization of the EMC and its role in *T. brucei* have remained unexplored. Interestingly, among protozoan parasites such as *Giardia intestinalis* and *Trichomonas vaginalis*, trypanosomes are the only organisms displaying a full set of (predicted) EMC subunits^1^. The conserved nature of the EMCs in conjunction with the position of *T. brucei* in the eukaryotic evolutionary tree provides a unique opportunity to study the role of the EMC in a highly diverged eukaryote that is essentially unrelated to Opisthokonts. This may help to reveal its ancestral function(s) in the ER and shed light on the root of its pleiotropy.

*T. brucei* is a eukaryotic protozoan parasite that causes African sleeping sickness in humans and nagana in cattle. Trypanosomes are digenetic parasites and have the ability to metabolically as well as morphologically adapt to vastly different environments in their insect vector and mammalian hosts^19,20^. Unlike mammals and yeast, trypanosomes have a single continuous mitochondrion harboring the mitochondrial genome or kinetoplast DNA^21,22^; its replication is in concert with the nuclear cell cycle, a peculiarity rarely seen in other eukaryotes^23,24^. *T. brucei* parasites are capable of *de novo* lipid synthesis and contain phospholipid compositions similar to other eukaryotes^25,26^, with phosphatidylcholine (PC) and phosphatidylethanolamine (PE) representing the two most abundant phospholipid classes^25,27^. In *T. brucei* bloodstream and procyclic forms, PC and PE are synthesized exclusively by the CDP-choline and CDP-ethanolamine, respectively, branches of the Kennedy pathway^25,28–30^.

In this study, we use biochemical and microscopic imaging approaches to identify the EMC in *T. brucei* (TbEMC) and to study possible functions of individual TbEMC subunits. Our work reveals that the TbEMC is composed of 9 subunits that are present in a complex localizing to the mitochondrial-ER interface. Interestingly, depletion of individual TbEMC subunits lead to selective disruption of *de novo* formation of PC or PE, re-emphasizing the role of the EMC in phospholipid synthesis.

## Results

### BLAST searches reveal homologs of human and yeast EMC subunits in *T. brucei*

By conducting pBLAST searches using the amino acid sequences of the individual members of the human and *S. cerevisiae* EMCs as queries, we identified 9 putative *T. brucei* EMC (TbEMC) proteins, with similarities of 20-40% with the corresponding yeast and human homologs (Fig. 1). This is in agreement with a previous phylogenetic analysis of EMC distribution in eukaryotes^1^. All TbEMC subunits except TbEMC2 and TbEMC8 are predicted to contain 1-3 TMDs, similar to their human and *S. cerevisiae* EMC homologs^15,31^ (Fig. 1), and TbEMC1, TbEMC3, TbEMC5, TbEMC7 and TbEMC10 are predicted to contain two signal peptides each, but according to iPSORT^32^ are not targeted to the mitochondrion.

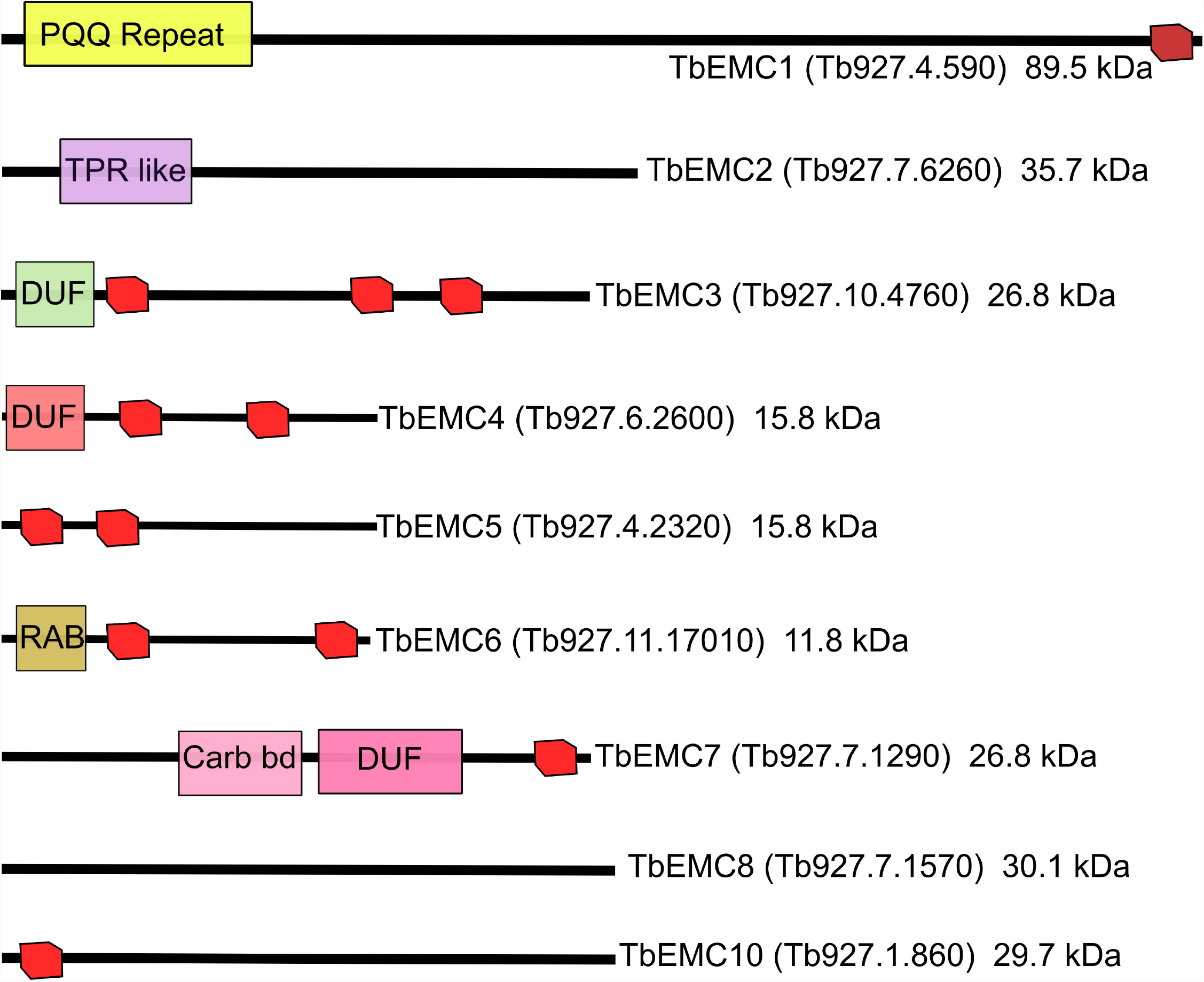

### Depletion of single TbEMC subunits inhibits *T. brucei* proliferation in culture

To study the essentiality of individual TbEMC subunits for the proliferation of *T. brucei* procyclic forms, we attempted to generate *null* mutants using CRISPR/CRISPR-associated protein 9-mediated gene deletion^33^. We successfully generated TbEMC7-knockout (TbEMC7-KO) and TbEMC10-KO cell lines and found that parasite proliferation was decreased after deletion of TbEMC7 (with an increase in cell doubling time from 9.0 h in parental to 12.5 h in TbEMC7-KO parasites) (Fig. 2A), but not after deletion of TbEMC10 (Fig. 2B). Deletion of the respective genes was verified by PCR (Fig. S1). Attempts to delete the genes encoding TbEMC1, TbEMC3, or TbEMC8 were not successful, which prompted us to down-regulate their expression using tetracycline-inducible RNA interference (RNAi). The results revealed that proliferation of parasites cultured in the presence of tetracycline to ablate TbEMC expression was arrested after approximately 2 days, demonstrating that expression of TbEMC1, TbEMC3 and TbEMC8 is essential for normal proliferation of *T. brucei* procyclic forms in culture (Fig. 2C-E; Fig. S2).

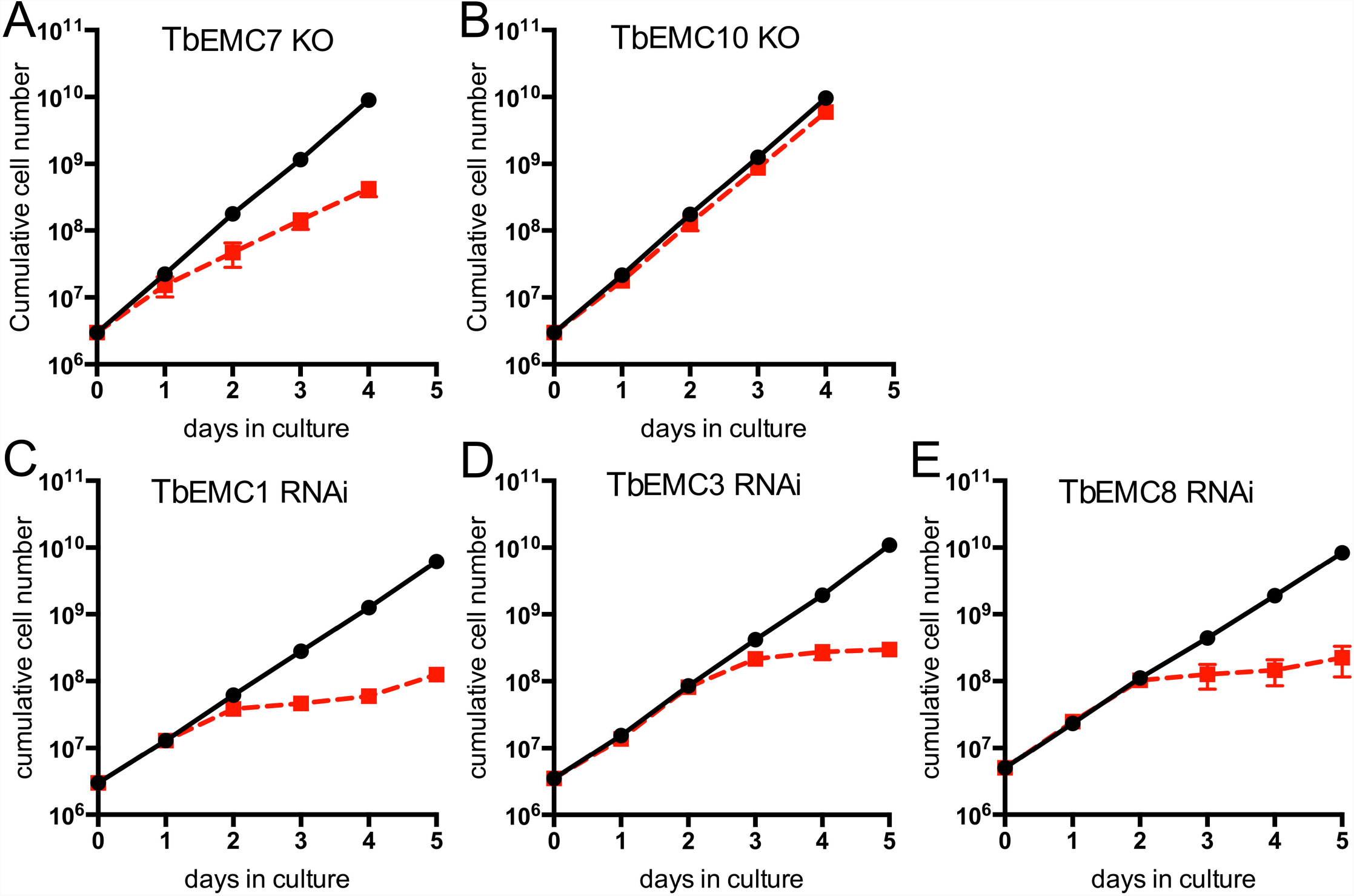

### Localization and organization of the TbEMC

To study their subcellular localization, we *in situ*-tagged all 9 TbEMC proteins in separate cell lines using HA (TbEMC1-4) or cMyc (TbEMC5-10) as tags. All TbEMCs were tagged C-terminally except TbEMC6, which was N-terminally tagged; C-terminal tagging of TbEMC6 was repeatedly unsuccessful. Analysis by SDS-PAGE and immunoblotting showed that all TbEMC proteins were expressed and migrated according to their predicted molecular masses (Fig. 3A, B). TbEMC8 and TbEMC10 consistently appeared as double bands. Analysis by immunofluorescence microscopy revealed that most TbEMC proteins co-localized with the ER lumenal marker protein BiP^34^, except TbEMC6 and TbEMC10, which exhibited preferential co-localization with ATOM40, the mitochondrial outer membrane protein-and tRNA-translocase in *T. brucei*^35,36^ (Fig. 4). At present, we cannot exclude that the N-terminal tag of TbEMC6 may affect its subcellular localization.

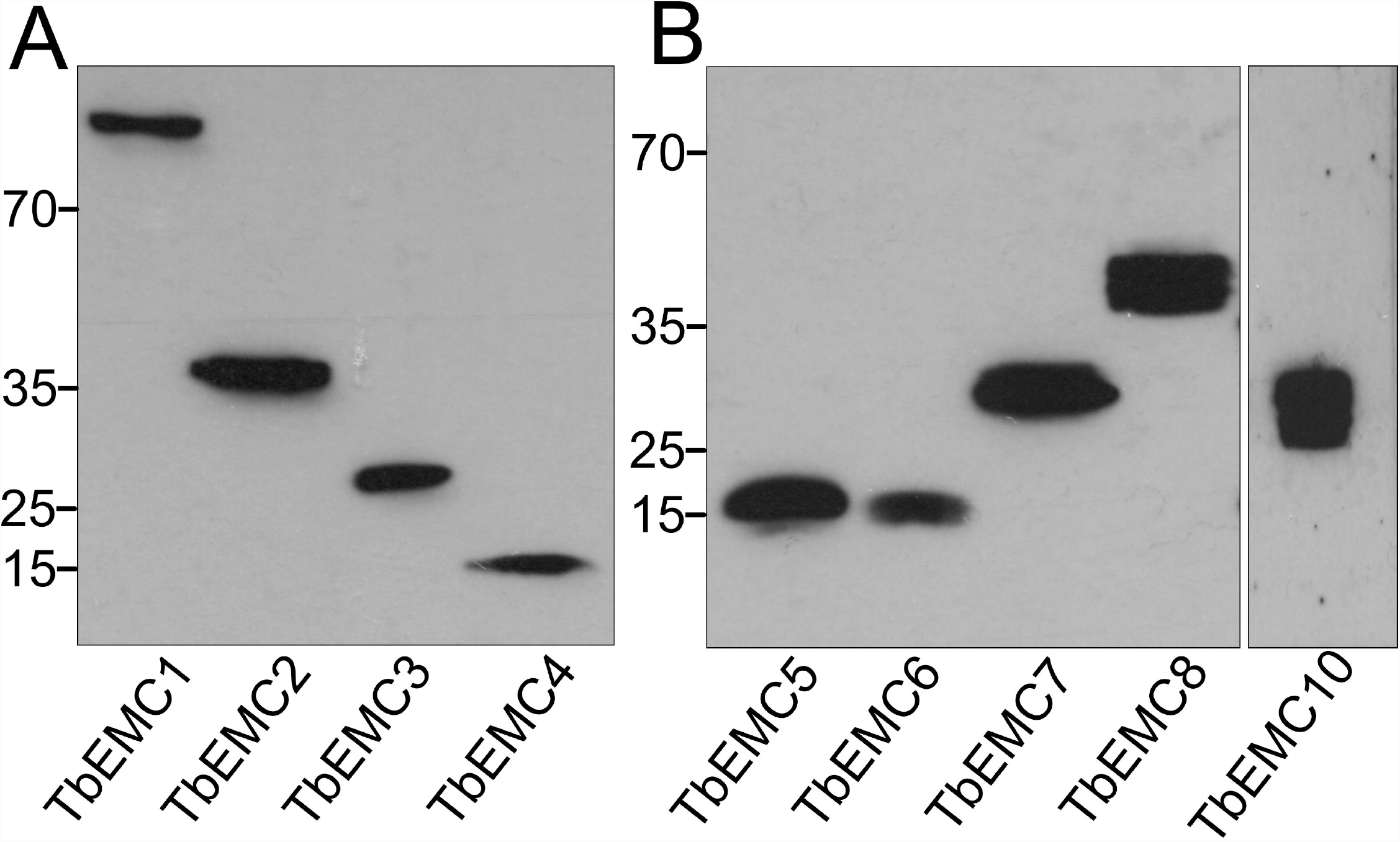

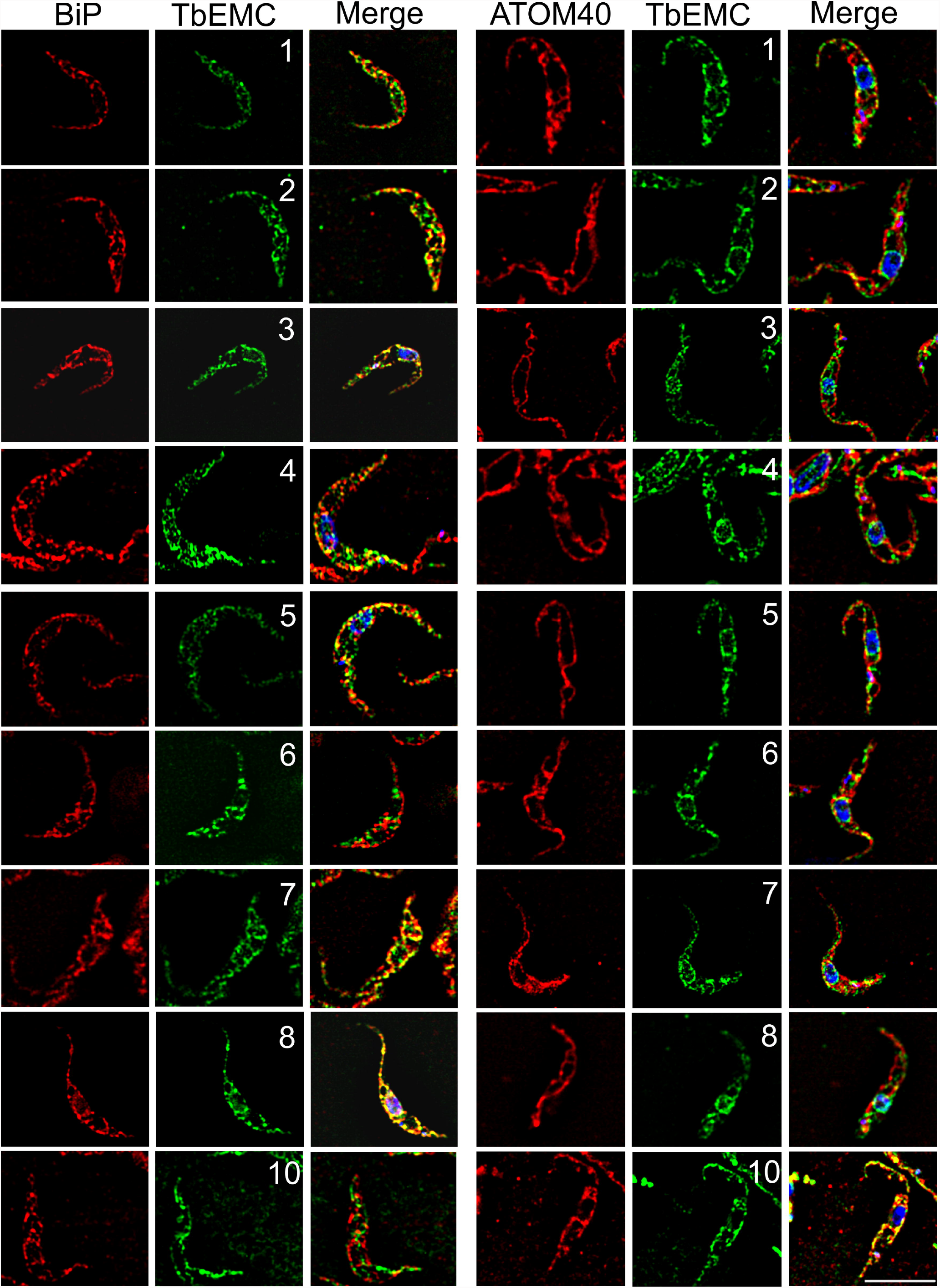

Most EMC subunits in mammalian cells and *S. cerevisiae* interact with each other^7,37^. To identify possible interactions between individual TbEMCs, we generated *T. brucei* procyclic forms co-expressing HA-and cMyc-tagged TbEMC proteins. We expressed TbEMC5-cMyc in tandem with TbEMC1-HA, TbEMC2-HA, TbEMC3-HA or TbEMC4-HA, and TbEMC3-HA in tandem with TbEMC6-cMyc, TbEMC7-cMyc, TbEMC8-cMyc or TbEMC10-cMyc, with each combination in separate parasites. Co-immunoprecipitation experiments were performed to probe for interactions between individual TbEMC proteins in the double-tagged parasites. Using TbEMC5-cMyc as bait, we were able to pull-down TbEMC1-HA, TbEMC2-HA, TbEMC3-HA and TbEMC4-HA (Fig. 5A-D). In reciprocal co-immunoprecipitation experiments, TbEMC5-cMyc was pulled-down only when using TbEMC2-HA as bait (Fig. 5A-D). In addition, when TbEMC3-HA was used as bait, TbEMC7-cMyc, TbEMC8-cMyc and TbEMC10-cMyc co-precipitated. TbEMC3-HA was faintly detected only in a pull-down using TbEMC8-cMyc as bait (Fig. 5E-G). TbEMC6-cMyc was not detected in the TbEMC3-HA or TbEMC8-cMyc precipitates.

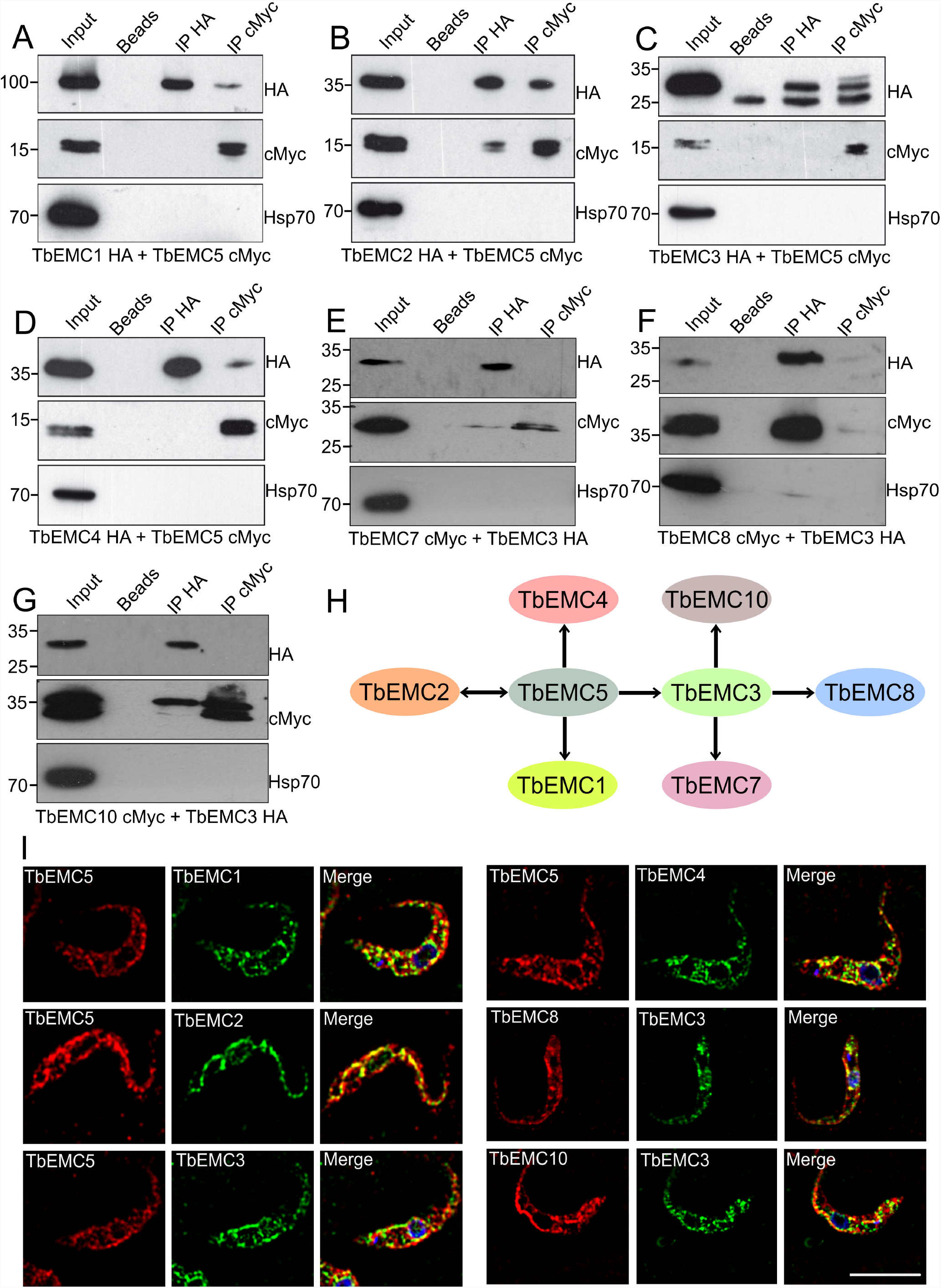

Furthermore, using immunofluorescence microscopy, we found high degrees of co-localization between TbEMC5-cMyc and TbEMC1-HA, TbEMC2-HA, TbEMC3-HA and TbEMC4-HA, while TbEMC3-HA and TbEMC8-cMyc showed a weaker co-localization (Fig. 5I). No co-localization was seen between TbEMC3-HA and TbEMC10-cMyc (Fig. 5I).

MS analyses of immunoprecipitates of TbEMC3-HA and TbEMC5-cMyc revealed the presence of several TbEMC proteins, further confirming physical interactions between individual subunits of the TbEMC. TbEMC3-HA co-precipitated TbEMC2, TbEMC8 and more than 20 ribosomal proteins of both the large and small ribosome subunits (Table S7). In addition, the pull-down was also enriched in glycosomal fructose-bisphosphate aldolase (Tb927.10.5620), tryparedoxin peroxidase (Tb927.9.5860), putative histone H2B (Tb927.10.10590), putative GTP binding nuclear protein RTB2 (Tb927.3.1120), mitochondrial carrier protein MCP12 (Tb927.10.12840) and nuclear RNA binding domain 2 (Tb927.11.14020). Furthermore, immunoprecipitation of TbEMC5-cMyc also co-precipitated several TbEMC subunits, including TbEMC8, TbEMC2, TbEMC3, TbEMC1 and TbEMC4. TbEMC5-cMyc also co-precipitated glycosomal membrane protein GIM5B (Tb927.9.11600) and the same putative GTP binding nuclear protein RTB2 (Tb927.3.1120) that was also pulled down by TbEMC3-HA (Table S8).

Together, our results show that all subunits of the TbEMC, except TbEMC6, individually interact with one or several other TbEMC proteins. In addition, the data indicates that the TbEMC is composed of TbEMC subunits only, i.e. it likely lacks additional tightly bound interaction partners. Although RTB2 (Tb927.3.1120) co-precipitated with both TbEMC3-HA and TbEMC8-cMyc, it is unlikely a component of the TbEMC as it localizes to the nuclear pore where it was found to mediate nucleocytoplasmic transport; its RNAi-induced depletion affected parasite growth and triggered programmed cell death^38^. Interestingly, the presence of a large number of ribosomal subunits in the TbEMC3-HA precipitate suggests association of the TbEMC with the ribosome. This would be consistent with the notion that the TbEMC might also be involved in co-translational protein insertion, as has been shown for human EMC^39^ and some substrates of its bacterial counterpart YidC^40^.

### TbEMC subunits form a high molecular mass complex

The yeast and mammalian EMC subunits form a single high molecular mass complex^2,7,16^. To investigate if the individual TbEMC proteins are also members of a complex in *T. brucei*, we performed native polyacrylamide gel electrophoresis (native PAGE) using *T. brucei* procyclic forms expressing the different epitope-tagged TbEMCs. Our results show that all 9 TbEMC proteins migrated as a high molecular mass complex of approximately 480 kDa after native PAGE analyses (Fig. 6A, B). An unspecific band at approximately 720 kDa appeared in all samples, including untagged parental parasites, when probed with the anti-HA antibody. Together with the TbEMC co-immunoprecipitation studies, these results demonstrate that the TbEMC subunits not only interact with each other but form a single high molecular mass complex.

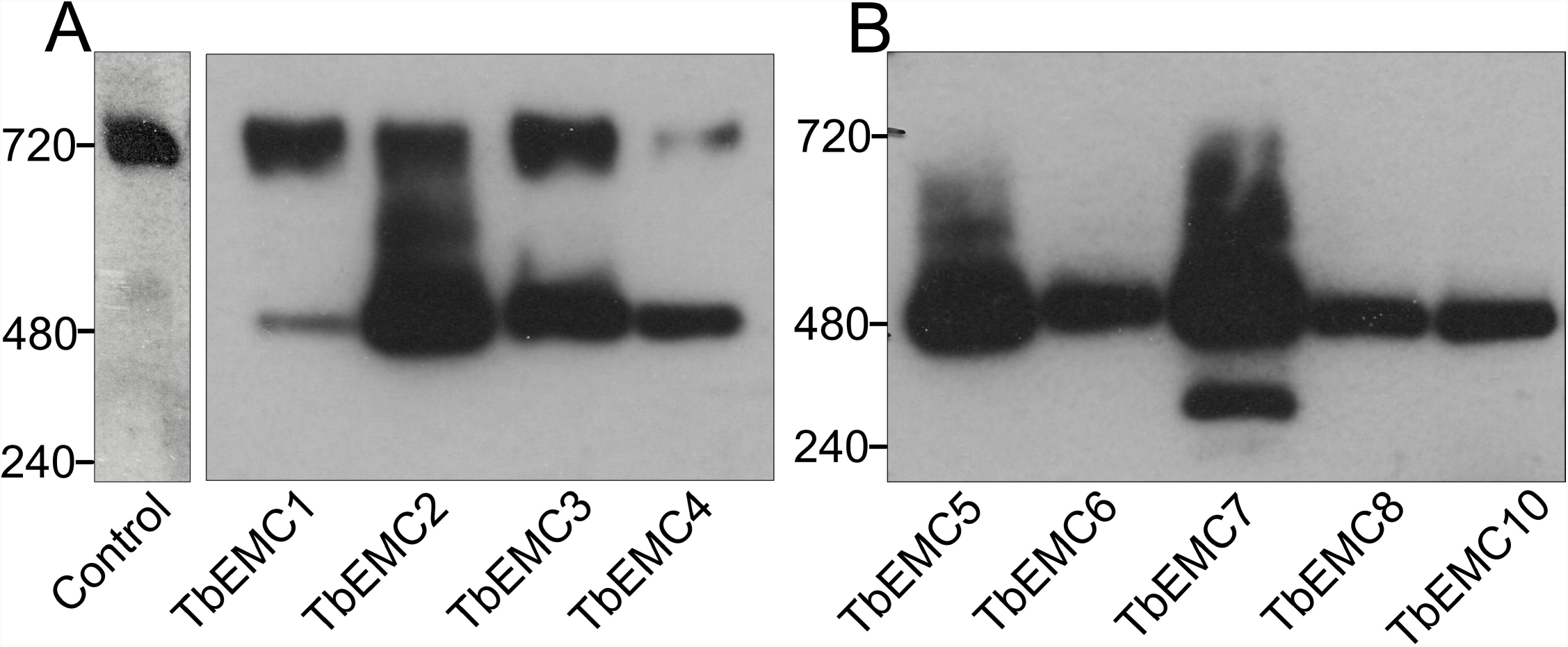

### Depletion of TbEMC7 destabilizes the TbEMC and leads to release of TbEMC3 from the complex

To study if the depletion of an individual subunit affects the integrity and/or stability of the TbEMC, we expressed *in situ*-tagged TbEMC3-cMyc in TbEMC7-KO parasites. Analysis by native PAGE revealed that in the absence of TbEMC7, TbEMC3-cMyc was not detectable (Fig. 7A), whereas in control parasites expressing TbEMC3-cMyc, the high molecular mass band at 480 kDa was clearly visible (compare also with Fig. 6A, showing parasites expressing TbEMC3-HA). Analysis of the same samples by immunoblotting under denaturing conditions demonstrated that TbEMC3-cMyc was expressed in the TbEMC7-KO background and that its expression level was similar to that in control parasites expressing TbEMC3-cMyc (Fig. 7B), ruling out the possibility that TbEMC3-cMyc itself is unstable in parasites lacking TbEMC7. These results suggest that in the absence of TbEMC7, TbEMC3-cMyc no longer associates with the complex and/or that the complex is destabilized. To distinguish between these possibilities, we expressed TbEMC8-cMyc in the TbEMC7-KO background and analyzed the presence of the TbEMC by native PAGE. The results show that in TbEMC7-KO parasites the 480 kDa band was reduced in intensity compared to control parasites (Fig. 7C), indicating that in the absence of the TbEMC7 subunit, the TbEMC is not properly formed or is destabilized. In line with these results, immunofluorescence microscopy analysis of TbEMC3-cMyc in TbEMC7-KO parasites revealed a diffuse staining pattern as compared to the punctate staining of TbEMC3-cMyc in control parasites (Fig. 7E), indicating that the TbEMC is not properly assembled. In contrast, in TbEMC10-KO parasites, the migration of TbEMC3-cMyc on native PAGE was unchanged compared to control cells (Fig. S3), indicating that the loss of TbEMC10 does not affect the integrity of the TbEMC.

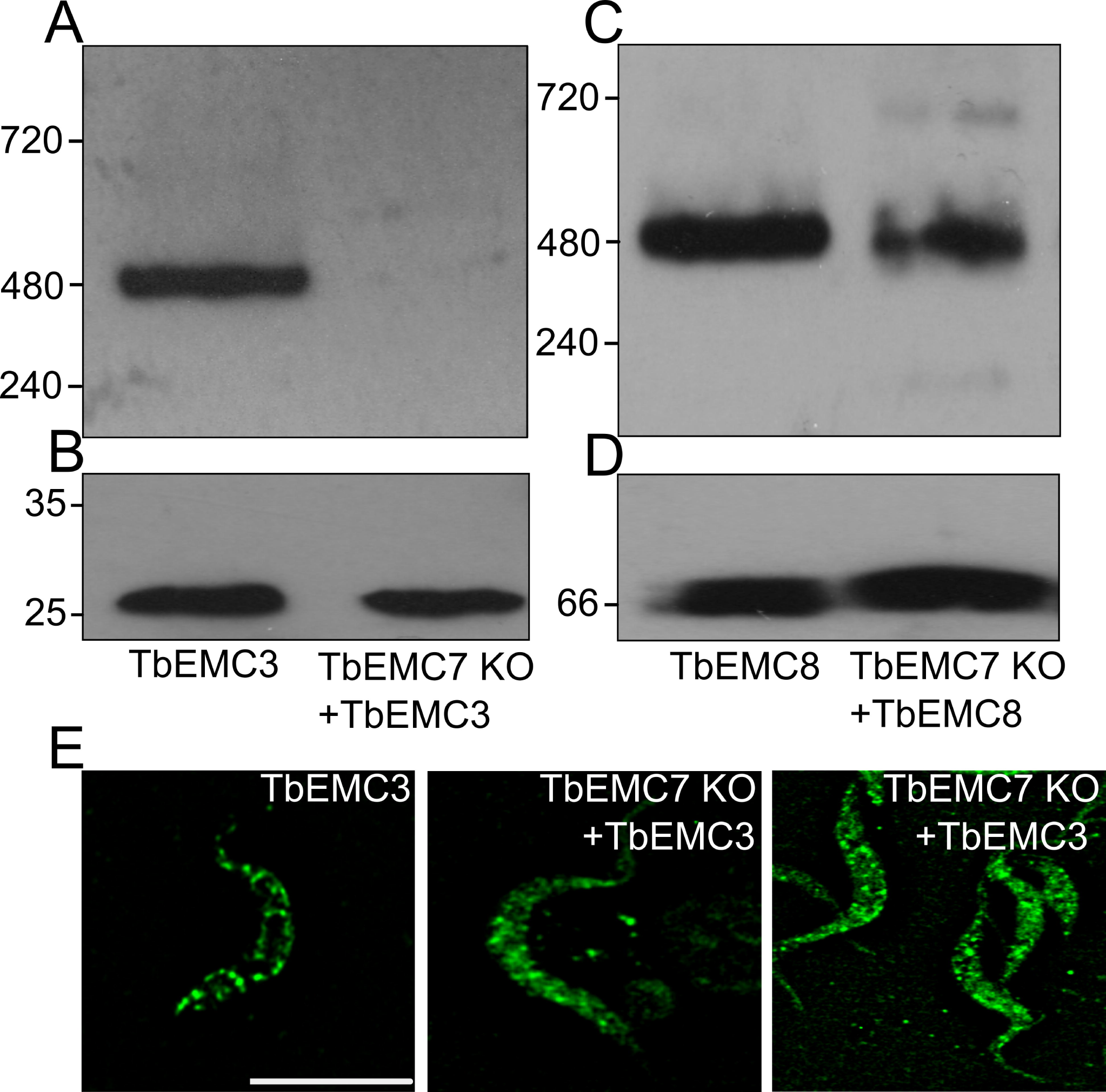

### The TbEMC localizes to the mitochondrial-ER interface

The EMC shows a considerable overlap with the ER and mitochondrion in *T. brucei*, and it is not unreasonable to assume that TbEMC clusters to the mitochondrial-ER interface, with the majority of the TbEMCs localizing to the ER and TbEMC6 and TbEMC10 interacting with the mitochondrial outer membrane (MOM). This would be in agreement with a previous study that implicates the EMC as an ER-mitochondrial tether in *S. cerevisiae*^7^. As conventional immunofluorescence microscopy techniques could not appreciably resolve the MOM and adjacent ER tubules, we wondered if the TbEMC could be traced biochemically to the mitochondrial ER-interface, in a separable membrane fraction termed mitochondria-associated ER membranes^41^. Established markers of these ER-subdomains associated with mitochondria are the phosphatidylserine synthases PSS1 and PSS2^42,43^. To establish the sub-organellar localization of PSS2 in *T. brucei*, we generated a cell line expressing TbPSS2 (Tb927.7.3760) containing a C-terminal cMyc-tag from an ectopic locus under the control of tetracycline and executed a well-established cell fractionation protocol to generate highly purified mitochondrial vesicles (Fig. 8A)^44^. The resultant fractions were analyzed by immunoblotting and probed for cMyc and the MOM marker ATOM40.

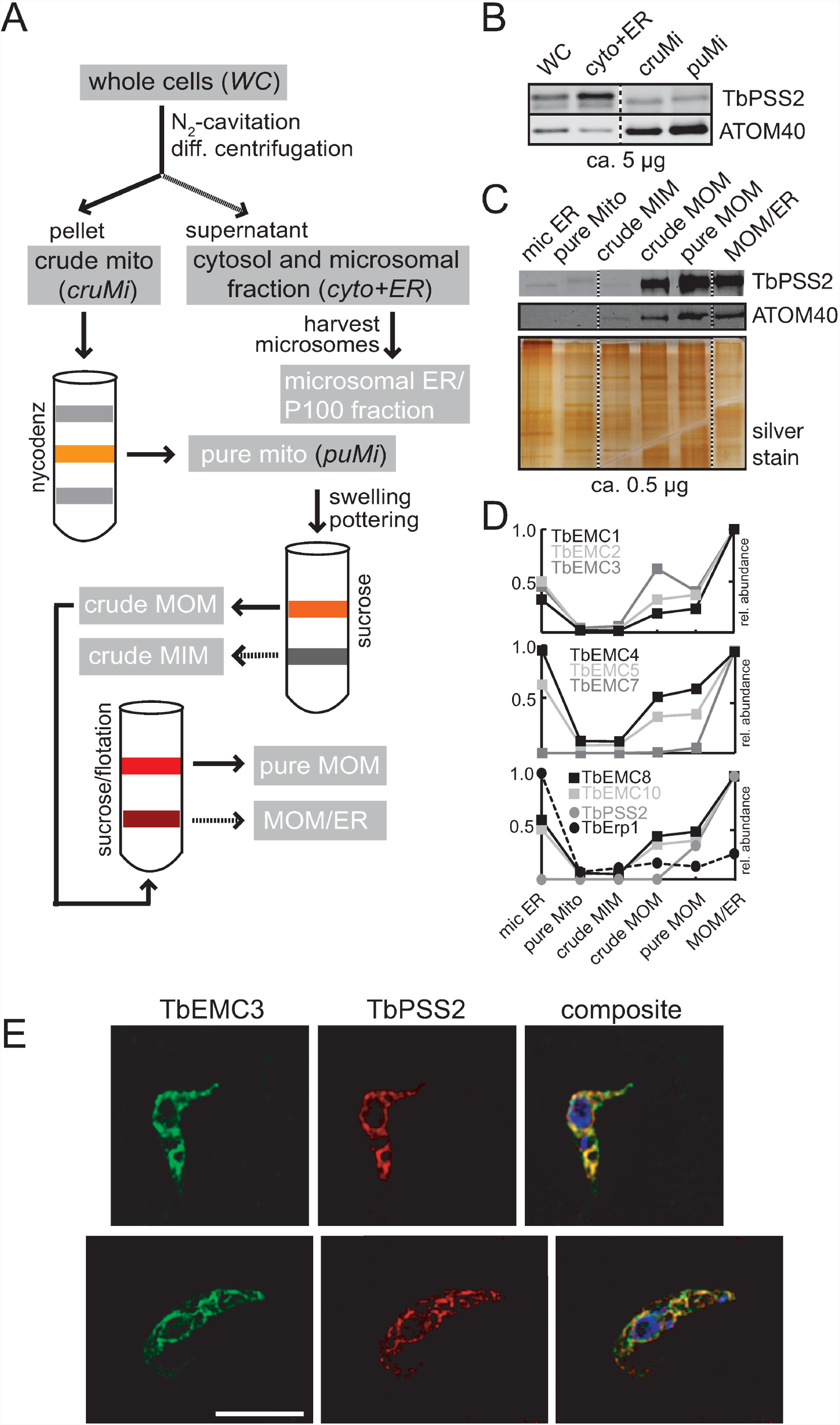

Our results show that while the majority of TbPSS2-cMyc is present in a fraction containing microsomal ER vesicles, a significant amount also co-fractionated with two mitochondrial fractions (Fig. 8B). Additionally, when we isolated the MOM from the purified mitochondria fraction (employing a previously established procedure^45^), TbPSS2-cMyc and ATOM40 were equally enriched in both the MOM and the fraction containing both MOM and ER proteins (MOM/ER fraction) (Fig. 8C). Thus, a fraction of TbPSS2-cMyc — despite being an ER resident protein^46^ — stably associated and co-fractionated with the MOM. We next asked if the TbEMC subunits would fractionate in similar fashion to TbPSS2-cMyc, indicative of a common localization at the mitochondrial ER-interface. To investigate this, we re-analyzed the data from our previous study in which the above-mentioned fractions (Fig. 8A) had been subjected to MS analyses to obtain protein abundance profiles^45^. We were able to confirm that all TbEMCs, except TbEMC6, were present in the individual fractions, with the protein abundance profiles shown in Fig. 8D, including the one for TbPSS2. Interestingly, we found that the protein abundance profiles of the TbEMCs closely matched that of TbPSS2, with maximal intensity in the MOM/ER fraction (Fig. 8D), whereas the ER-resident protein TbErp1^47^ shows maximal intensity in the microsomal ER fraction, serving as a negative control. These results are in good agreement with the immunoblot analyses shown in Fig. 8C.

Finally, to confirm co-localization of the TbEMCs with TbPSS2 by immunofluorescence microscopy, we generated a cell line co-expressing TbEMC3-HA and TbPSS2-cMyc. Our results show that TbEMC3-HA strongly co-localized with TbPSS2-cMyc (Fig. 8E). Together, these data indicate that the TbEMC proteins and TbPSS2 localize to the same sub-organellar structure, which we refer to here as the mitochondrial ER-interface. Inter-organellar contact sites between the ER and the mitochondrion have not been studied before in *T. brucei*.

### Identification of TbEMC substrates

A combined approach of RNAi and stable isotope labeling with amino acids in cell culture (SILAC)-based quantitative proteomics has previously been used to determine the substrates of the trypanosomal atypical mitochondrial outer membrane translocase (ATOM complex)^49^, and the two mitochondrial outer membrane protein import receptors ATOM46 and ATOM69^50^, respectively. In both cases it was sufficient to analyze whole cell extracts since non-imported mitochondrial proteins that accumulate in the cytosol are rapidly degraded. We therefore applied the same approach to identify the substrates of the TbEMC by performing SILAC-based proteomics analyses of whole cells after RNAi-mediated depletion of TbEMC3 and TbEMC8 for 3 and 3.5 days. The volcano plots in Fig. 9 show that, in both cell lines, the targets of the RNAi, either TbEMC3 or TbEMC8, were efficiently downregulated at both time points. Moreover, more than 3000 proteins were quantified in each data set (for information about proteins identified and quantified, see Table S9).

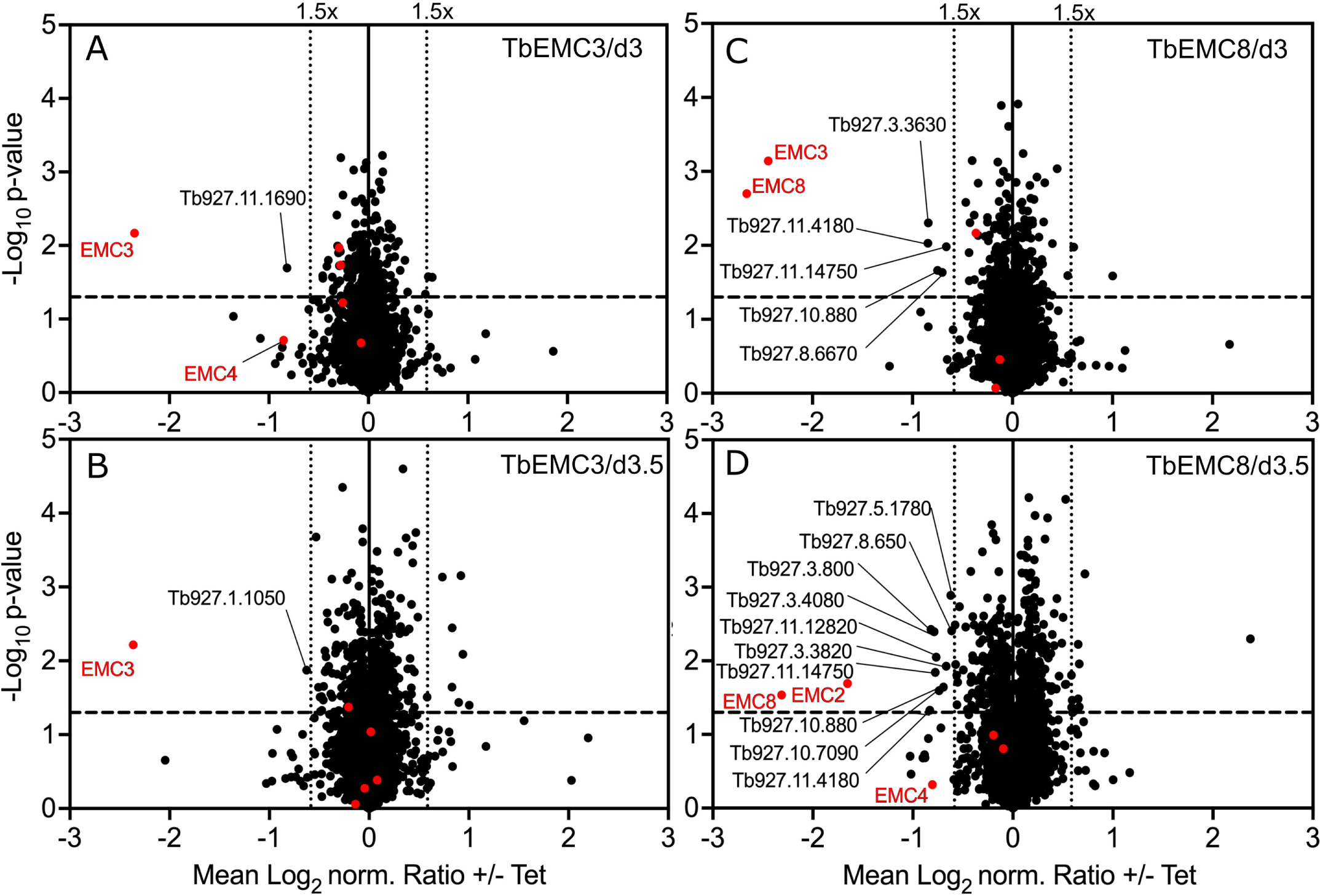

In the TbEMC3 RNAi cell line we did detect significant downregulation of TbEMC10 and TbEMC2 at at least one time point, however, the extent of downregulation was much lower, at approximately 1.2-fold. Interestingly, the only other proteins that were downregulated >1.5-fold at at least one of the time points were putative emoplamil binding protein (Tb927.11.1690) and metallo-dependent phosphatase-like domain-containing protein (Tb927.1.1050). Both proteins contain multiple predicted transmembrane domains and localize to the ER in human tissue (for the homolog of Tb927.11.1690)^51^ and *T. brucei* (Tb927.1.1050) (Fig. 9A, B). Thus, it is possible that these two proteins require the TbEMC for membrane insertion.

In addition, RNAi against TbEMC8 revealed significant downregulation of TbEMC3 (>5-fold) and TbEMC2 (>3-fold) (Fig. 9C, D). Each protein was detected at only one time point. Other TbEMC proteins were depleted to lesser extents, such as TbEMC1 and TbEMC4 (Fig. 9C, D). Furthermore, we noticed downregulation of >1.5-fold of 11 proteins at at least one of the time points. Five of them contain predicted transmembrane domains and, therefore, with the exception of mitochondrial alternative oxidase AOX (Tb927.10.7090), are at least initially inserted into the ER membrane indicating they could be possible substrates of the TbEMC. However, we also detected predicted soluble proteins that would not be inserted into the ER membrane, suggesting that some of the downregulated proteins are likely affected indirectly by TbEMC8 depletion.

Together, the SILAC-based RNAi analyses confirm a mutual dependence between certain TbEMC subunits that could be expected from proteins within a stable complex. However, they also show that we can only detect few putative substrates of the TbEMC using this method. A possible explanation for this could be that in contrast to what was observed with mitochondria, non-inserted TbEMC substrates may not be immediately degraded in the cytosol. Furthermore, it is possible that if TbEMC substrates are not inserted into the ER membrane, they might be mislocalized to other membranes. In either case we would not detect them with our approach.

### Depletion of individual TbEMC proteins inhibits *de novo* phospholipid synthesis

Besides its role as membrane protein insertase, the EMC has also been proposed to play a role in phospholipid homeostasis. To study if the TbEMC may play such a role in *T. brucei*, we analyzed *de novo* synthesis of the major phospholipid classes in *T. brucei* procyclic forms by measuring incorporation of [^3^H]-choline and [^3^H]-ethanolamine into PC and PE, respectively, after depletion of individual TbEMC subunits. The labeled precursors were added after 48 h of RNAi, i.e. before parasite growth was affected by down-regulation of the individual TbEMC proteins (see Fig. 2C-E), and incubated for 4 h to allow uptake and metabolism into phospholipids via the Kennedy pathway^30^. Formation of [^3^H]-PI using [^3^H]-*myo*-inositol as precursor was used as internal labeling control. Lipids were separated by thin layer chromatography (TLC) and incorporation of radioactivity into different phospholipid classes was quantified by radioisotope scanning; a representative TLC analysis is shown in Fig. S4. Interestingly, we found that depletion of TbEMC1 or TbEMC3 decreased *de novo* synthesis of [^3^H]-PC by 50-60% compared to control parasites (Fig. 10A, C). In contrast, down-regulation of neither of these two TbEMCs had an effect on the synthesis of [^3^H]-PE (Fig. 10B, D). Unexpectedly, the opposite effect was observed after depletion of TbEMC8. RNAi against TbEMC8 decreased *de novo* synthesis of [^3^H]-PE by >60% compared to control parasites, while no effect was seen for [^3^H]-PC formation (Fig. 10E, F). No major differences in [^3^H]-PC or [^3^H]-PE synthesis was seen in TbEMC7-KO (Fig. 10G, H) and TbEMC10-KO parasites (Fig. 10I, J). Together these results show that in *T. brucei* procyclic forms depletion of individual TbEMC proteins results in disturbance of select pathways for *de novo* synthesis of the major phospholipid classes.

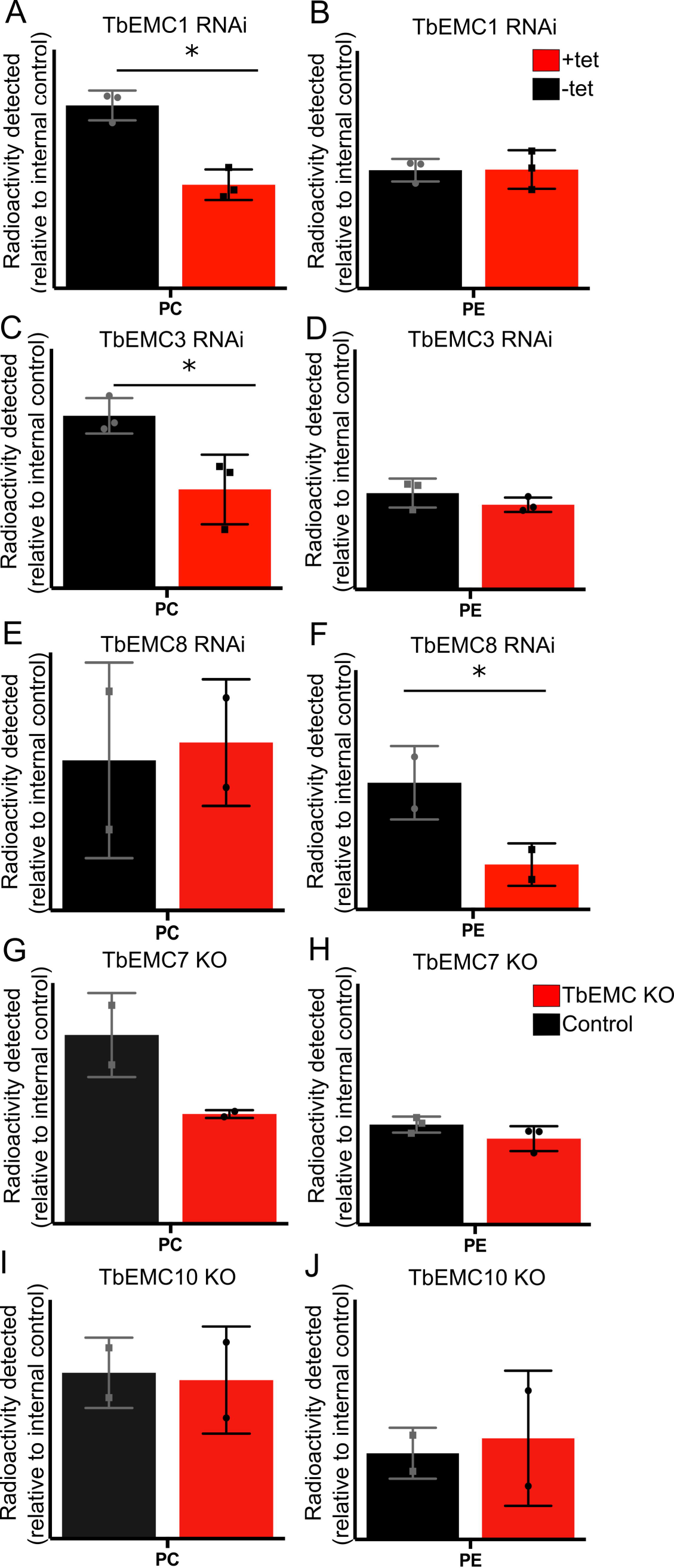

However, these effects cannot be related to depletion of individual enzymes of the Kennedy pathway. Several of these enzymes, including choline/ethanolamine kinase (Tb927.5.1140), choline kinase (Tb927.11.2090), ethanolamine-phosphate cytidylyltransferase (Tb927.11.14140), choline-phosphate cytidylyltransferase (Tb927.10.12810) and ethanolamine phosphotransferase (TbEPT; Tb927.10.13290), were identified in the SILAC-based RNAi experiments, however, none of them were significantly downregulated in response to TbEMC8 or TbEMC3 depletion (Fig. 9). Choline/ethanolamine phosphotransferase (TbCEPT; Tb927.10.8900) was not detected. These data suggest that the observed effects of TbEMC depletion on PC and PE biosynthesis are not caused by reduced levels of enzymes of the Kennedy pathway. The data however do not exclude the possibility that the two membrane-bound enzymes, TbEPT and TbCEPT, were re-localized to other membranes during TbEMC depletion.

## Discussion

In humans, EMC1, EMC2, EMC3, EMC5 and EMC6 form the core of the complex while EMC4, EMC7 and EMC10 constitute peripheral components. In addition, EMC8 and EMC2 form an interface with EMC5^15^. In yeast, all EMC subunits interact with each other, except EMC1-EMC3 and EMC1-EMC4^7^. In the present study we show that all individual TbEMC subunits interact with at least one other subunit, except TbEMC6, which did not interact with either TbEMC3 or TbEMC8 in our pull-downs or in the MS-based co-immunoprecipitation analyses. Interestingly, unlike in mammalian cells where EMC7 is considered a peripheral component of the complex^16^, we identified TbEMC7 as a crucial subunit integral to the stability of the TbEMC. In its absence, the TbEMC is not properly formed or is destabilized, resulting in at least the release of TbEMC3 from the complex. TbEMC7-KO parasites continue to synthesize PC, albeit less efficiently, and are viable. In contrast, RNAi-mediated depletion of TbEMC3 results in growth arrest. Together, these results indicate that TbEMC3 is able to, at least partially, maintain its role in PC synthesis even after release from the TbEMC. Alternatively, TbEMC3 may retain its function in PC synthesis via transient association with the TbEMC in TbEMC7-KO parasites.

The EMC in mammalian and *S. cerevisiae* cells localizes exclusively to the ER. Analogously, most TbEMC subunits also localized to the ER and formed a complex in *T. brucei*. However, unexpectedly, TbEMC6 and TbEMC10 exhibited mitochondrial localization. This, together with our biochemical data showing that the TbEMC proteins and TbPSS2 co-fractionate in a distinct sub-organellar membrane fraction, localizes the TbEMC to the mitochondria-ER interface. A similar localization of the EMC in close proximity to both organelles has been proposed in *S. cerevisiae*, where it was postulated to act as an ER-mitochondrial tether^7^.

Besides its function as ER membrane protein insertase, the EMC has been implicated in inter-organellar lipid exchange^7,10^. Although lipidomic analyses have revealed that the loss of EMC3 leads to perturbations in cellular lipid levels in murine cells^10^, involvement of individual EMC subunits in the intricate pathways of lipid biosynthesis has not been studied before in detail. Here we demonstrate that RNAi-mediated ablation of TbEMC1, TbEMC3 or TbEMC8 expression inhibited production of PC or PE, the major phospholipid classes in *T. brucei* procyclic forms. PC and PE in *T. brucei* parasites are synthesized exclusively by the CDP-choline and CDP-ethanolamine branches of the Kennedy pathway^26^, respectively. Both branches are essential for parasite survival in culture. The first two reactions of either branch, catalyzed by the respective kinases and phosphotransferases, occur in the cytosol. In contrast, the final reactions are mediated by the membrane-bound enzymes TbCEPT and TbEPT, respectively, in the ER. The two enzymes localize to different sub-compartments of the ER, with TbCEPT being present exclusively in the perinuclear ER and TbEPT localizing mostly to the bulk ER, with a fractional overlap with the perinuclear ER^30^. Since both enzymes have multiple TMDs, we hypothesized that their expression and/or membrane insertion might be affected following disruption of the TbEMC’s attributed main function as membrane protein insertase. However, our SILAC data provided no evidence for downregulation of TbEPT or TbCEPT after the induction of RNAi against TbEMC proteins. In addition, since down-regulation of TbEMC1 or TbEMC3 resulted in inhibition of *de novo* production of PC, but had no effect on PE synthesis, while depletion of TbEMC8 had the opposite effect, i.e. inhibited *de novo* formation of PE without affecting PC synthesis, the effects cannot be explained by a general defect in membrane protein insertion. The localization of the TbEMC to the mitochondria-ER interface may suggest that certain subunits could be involved in the transfer of phospholipid precursors or intermediates between these organelles, or that the phospholipid synthesis defect was caused by disruption of this interface. Alternatively, recent work has shown that the presence of a hydrophilic vestibule or substrate binding pocket is central to the insertion mechanism of the EMC complex^15,31^. Simulations on the *S. cerevisiae* EMC structure have revealed that perturbations in the cytoplasmic leaflet of the ER bilayer facilitates flooding of the substrate binding pocket with water, promoting a hydrophilic environment instrumental to insertion^52^. Since PC and PE are major structural components of the ER membrane, it is tempting to speculate that the role of the TbEMC in lipid biosynthesis may be tied to the creation of a microenvironment conducive to TMD insertion, and thus TbEMC function.

In an attempt to identify substrates that may be inserted into the ER membrane by the TbEMC, we performed SILAC and RNAi experiments using TbEMC3 and TbEMC8 as RNAi targets. Technically these experiments worked well, as demonstrated by the efficient downregulation of the target proteins. However, the levels of only few other proteins were significantly downregulated more than 1.5-fold. Among these only few are predicted ER membrane proteins which could represent substrates of the TbEMC acting as insertase. However, the pleiotropic effects of TbEMC depletion in *T. brucei* (or the EMC in other organisms) suggest that the complex has more than a handful of substrates. This raises the question of why we were unable to detect the substrates in our SILAC-based approach. One reason may be that we only analyzed whole cell proteomes and that, unlike for mitochondria, where non-imported proteins are rapidly degraded in the cytosol, non-inserted TbEMC substrates are not degraded. It is possible that in the absence of the TbEMC, certain TbEMC substrates are inserted into the ER by other means (e.g. via the Sec61 pathway), or that non-inserted substrates accumulate as stable aggregates in the cytosol. Furthermore, we cannot exclude that failed insertion of TbEMC substrates into the ER membrane leads to their mislocalization to other membranes. In all these cases, the total cellular levels of the TbEMC substrates would not change upon TbEMC depletion. Further work analyzing the proteomes of subcellular fractions may help to distinguish between these different possibilities but are beyond the scope of the present work.

## Materials and Methods

### Reagents

All reagents were of analytical grade and purchased from Merck-Sigma Aldrich (Darmstadt, Germany) or MilliporeSigma (Burlington, MA, USA) unless otherwise stated. Kits for plasmid DNA extractions, PCR purifications, vector dephosphorylation and ligation reactions were purchased from Promega (Wallisellen, Switzerland). All sequencing was done by Microsynth AG (Balgach, Switzerland).

### Cell culture and knockdown trypanosome cell lines

Procyclic form trypanosomes were cultured at 27 °C in SDM79^53^ supplemented with 10% heat inactived fetal bovine serum (Thermo Fisher Scientific, Reinach, Switzerland) and a mixture of hemin and folic acid (160 µM hemin + 90 µM folic acid). Procyclic form (SmOxP9) trypanosomes expressing tagged TbEMCs were cultured in the presence of hygromycin (25 μg/ml), G418 (15 μg/ml), puromycin (2 μg/ml), blasticidin (5 μg/ml). TbEMC-KO parasites were cultured in the presence of hygromycin (25 μg/ml), G418 (15 μg/ml), puromycin (2 μg/ml) and blasticidin (5 μg/ml), and TbEMC-KO parasites expressing in-situ tagged proteins were cultured in the presence of an additional 0.2 μg/ml of phleomycin. *T. brucei* strain 29-13^54^ was used to generate RNAi cell lines against TbEMC1, TbEMC3 and TbEMC8, and cultured in the presence of hygromycin (25 μg/ml) and G418 (15 μg/ml), and puromycin (2 μg/ml), blasticidin (5 μg/ml) and phleomycin (0.2 μg/ml) in TbEMC1 RNAi, TbEMC3 RNAi and TbEMC8 RNAi, respectively, parasites. For induction of RNAi tetracycline (1 μg/ml) was added. The RNAi constructs were based on a derivate of the stem-loop vector pLEW100^54^ using the following inserts for the RNAi construct: TbEMC1 (Tb927.4.590) ORF nt 996-1246, TbEMC3 (Tb927.10.4760) 3’ UTR nt 13-165 and TbEMC8 (Tb927.8.1570) ORF nt 2084-2501. Transfection and selection of clones of the above-mentioned TbEMC knock-downs were performed as described previously^55^ and RNAi efficiency was verified by mRNA Northern blotting.

### Generation of TbEMC-KO parasites using clustered regularly interspaced short palindromic repeats (CRISPR)/CRISPR associated protein 9

CRISPR/Cas9 was employed to create knockouts of individual TbEMCs, as described previously^33^. Briefly, procyclic form (SmOxP9) trypanosomes were used to create knockouts by replacing the TbEMC gene on both alleles with 2 antibiotic resistance cassettes by transfection. The 2 antibiotic resistance cassettes flanked by homology sequences of 30 nt were amplified by PCR reactions using pPOTv6 plasmids^56^ with hygromycin and geneticin resistance cassettes. Both the antibiotic cassettes as well as short guide RNA templates consisting of a T7 promoter sequence, a Cas9 binding site, and 20 nt long target sequence were amplified as described previously^33^. Primers were generated by www.leishgedit.net (Table S5) and all PCR reactions were performed using Expand High Fidelity PCR System (Roche, Basel, Switzerland), pooled and purified using PCR purification kits. Transfection was performed using a 4DLNucleofector system, with the FI-115 program (Lonza, Basel, Switzerland). The transfected culture was distributed to 24Lwell plates after a 2 h recovery and 22.5 μg/ml geneticin and 37.5 μg/ml hygromycin (selective antibiotics) were added after 24 h.

TbEMC-KO genotypes were verified by PCR analysis of genomic DNA isolated from parental and knockout cell lines. All primer pairs and PCRs conducted for TbEMC knockout verification are listed in Fig. S1.

### In-situ tagging of TbEMCs

TbEMCs were tagged in SmOxP9 as well as TbEMC-KO parasites using CRISPR/Cas9 technique as described above. Briefly, resistance cassette consisting of a cMyc tag sequence and a blasticidin/hygromycin/phleomycin resistance gene and homology sequences was inserted in TbEMC genes by homologous recombination. Corresponding guide RNA templates were also generated. All primers were generated by www.leishgedit.net (Table S6). The PCR products were pooled, purified, and transfected as described above. 24 hours after distribution into 24-well plates, selection antibiotics were added. TbEMC protein expression in the parasites was analyzed by SDS-PAGE and immunoblotting. TbPSS2 cell line has been described previously^46^.

### Whole cell lysates, crude membrane preparations and immunoprecipitations

For whole cell lysate preparations, 10^7^ trypanosomes were harvested, and the resultant pellet was washed with Tris-buffered saline (TBS, 10 mM Tris-HCl, pH 7.5, 144 mM NaCl) and resuspended in sample buffer (2.5% SDS). The sample was then incubated at 50 °C for 5 min.

For crude membrane preparations, a digitonin extraction was performed. Trypanosomes (10^8^ cells) were harvested by centrifugation, washed in TBS and resuspended in 0.5 ml buffer (20 mM Tris-HCl, pH 7.5, 600 mM sorbitol, 2 mM EDTA) containing 0.05% (w/v) digitonin. The suspension was centrifuged at 6000 x g. Subsequently, 100 μl extraction buffer (20 mM TrisLHCl, pH 7.2, 15 mM NaH_2_PO_4_, 0.6 M sorbitol) containing 1.5% (w/v) digitonin was added to the pellet. The sample was incubated on ice for 15 min and membranes were pelleted by centrifugation at 16’000 x g in a tabletop centrifuge (15 min, 4 °C). The resultant pellet was used for further analysis.

For immunoprecipitations, whole cell lysates were solubilized in 1 ml lysis buffer (10 mM Tris-HCl, pH 7.4, 150 mM NaCl, 1 mM EDTA, 1% Triton X-100, 0.5% NP-40, protease inhibitors) and incubated on ice for 30 min. After centrifugation at 16’000 x g (15 min, 4 °C) the lysate (input) was incubated with anti-HA or anti-cMyc antibodies (9E10; Santa Cruz Biotechnology, Dallas, TX, USA / 16B12 Enzo Life Sciences, Farmingdale NY, USA; dilution 1:40) in combination with 50 μl Protein G Dynabeads (Thermo Fischer Scientific) for 16 h. Bound proteins were eluted by addition of sample buffer and heating at 50 °C for 5 min. The proteins were analyzed by SDS-PAGE and immunoblotting.

Immunoprecipitation followed by MS analysis was done by washing 10^8^ parasites in PBS, followed by cross-linking in 0.1% paraformaldehyde in PBS for 8 min at room temperature. After addition of glycine to a final concentration of 125 mM for 5 min, cells were washed twice with TBS. Cell pellets were lysed and processed as described above, except that the beads were washed with lysis buffer lacking detergent after the immunoprecipitation. Washed beads were frozen in liquid nitrogen and stored at -80 °C.

### PAGE

Proteins were denatured by SDS and run on 10% polyacrylamide gels as described previously^57^. For native PAGE, samples were prepared and loaded onto 4-12% gradient gels as described elsewhere^58^. The proteins were then transferred onto PVDF membranes (Thermo Fisher Scientific) using a semidry protein blotting system (Bio-Rad, Hercules, CA, USA) for 75 min at a maximal current of 2.5 mA cm^−2^ gel area. The membranes were then blocked for 1 h in TBS, containing 5% (w/v) milk powder, and proteins of interest were identified using specific primary and horseradish peroxidase-conjugated secondary antibodies. Protein-specific antibodies were visualized using SuperSignal™ West Pico PLUS Chemiluminescent Substrate (Thermo Fisher Scientific). The following antibodies were diluted in TBS, containing 5% milk: mouse monoclonal anti-cMyc (9E10; Santa Cruz Biotechnology; dilution 1:1000); mouse monoclonal anti-HA (HA11; Enzo Life Sciences; dilution 1:3000); rabbit anti-AAC1 (kindly provided by Alena P. Ziková, University of South Bohemia, Ceske Budejovice, Czech Republic; dilution 1:20000); HRP-conjugated anti-rabbit and anti-mouse antibody (DAKO-Agilent, Basel, Switzerland; dilution 1:1000 and 1:5000, respectively).

### Immunofluorescence microscopy

For immunofluorescence staining of trypanosomes, 10^6^ cells were harvested by centrifugation, resuspended in ice cold PBS (phosphate-buffered saline; 137 mM NaCl, 2.7 mM KCl, 10 mM Na_2_HPO_4_, 2 mM KH_2_PO_4_, pH 7.4), allowed to adhere onto glass slides (Thermo Fisher Scientific) and fixed with 4% (w/v) paraformaldehyde for 10 min. The cells were then washed with PBS and permeabilized with 0.2% (w/v) Triton X-100 for 5 min. After blocking in PBS containing 2% (w/v) BSA, for 30 min, parasites were incubated with antibodies diluted in blocking solution for 45 min. The following primary antibodies were used: rabbit anti-cMyc (LubioSciences, Zurich, Switzerland; dilution 1:200); mouse monoclonal anti-cMyc (9E10, Santa Cruz Biotechnology; dilution 1:200), mouse anti-HA (16B12; Enzo Life Sciences; dilution 1:200), rabbit anti-ATOM40 (dilution 1:1000), rabbit anti-BiP (kindly provided by J. Bangs, University of Buffalo, NY, USA; dilution 1:20,000). After 3 washes in PBS for 5 min each, the cells were incubated with the corresponding secondary fluorophore-conjugated antibodies Alexa Fluor goat anti-mouse 488 and goat anti-rabbit 594 (Thermo Fisher Scientific; dilution 1:1000) in blocking solution for 45 min. Cells were washed, air dried and mounted with Vectashield (Vector Laboratories, Burlingame, CA, USA) containing 4`,6-diamidino-2-phenylindole (DAPI).

Fluorescence microscopy was performed with a Leica DM 16000 B inverted microscope using a 60x oil objective and images were acquired using Leica DFC360 FX camera. Image deconvolution and processing was performed using Leica LAS X and Fiji software (NIH).

### [^3^H]-glycerol labeling and lipid analysis

Trypanosomes (10^8^ cells) were labeled with 50 μCi [^3^H]-ethanolamine or 50 μCi [^3^H]-choline chloride in combination with 20 μCi [^3^H]-inositol (Anawa, Kloten, Switzerland) for 4 h before being harvested by centrifugation and washed twice with TBS. The radiolabeled lipids were extracted according to Bligh and Dyer^59^, spotted onto silica gel plates (Merck, Zug, Switzerland) and separated by one dimensional TLC using a solvent system consisting of chloroform: methanol: acetic acid (65: 25: 8, by vol) to track PE formation and chloroform: methanol: acetic acid: water (50: 30 : 8: 4, by vol) to track PC formation. The plates were dried, and radioactivity was detected using a radioisotope scanner (Berthold Technologies, Bad Wildbad, Germany). Data processing was done using the Rita Control software provided by the manufacturer.

### Preparation of mitochondrial vesicles, mitochondrial outer membrane (OM) and OM/ER fractions

The detailed procedure for the purification of mitochondrial vesicles from a variety of different cell amounts is detailed eslewhere^44^. Briefly, trypanosomes were lysed at isotonic conditions using nitrogen cavitation, and following differential centrifugation, mitochondrial vesicles were purified from Nycodenz step-gradients subjected to density gradient centrifugation. Mitochondrial OM was further purified from these mitochondrial vesicles according to a previously described protocol^45^.

### SILAC analysis

Cells were washed in PBS and taken up in SDM-80 supplemented with 5.55 mM glucose, either light (^12^C_6_/^14^N_χ_) or heavy (^13^C_6_/^15^N_χ_) isotopes of arginine (1.1 mM) and lysine (0.4 mM) (Euroisotop, Saarbrücken, Germany) and 10% dialyzed fetal calf serum (BioConcept, Allschwil, Switzerland). To guarantee complete labeling of all proteins with heavy amino acids, the cells were cultured in SILAC medium for 6–10 doubling times. The TbEMC3 and TbEMC8 RNAi cell lines were induced for 3 or 3.5 days, and 1×10^8^ uninduced and 1×10^8^ induced cells were harvested and mixed. The pellets generated from mixing these cells were then analyzed. All SILAC-based RNAi experiments were performed in three biological replicates including a label-switch and analyzed by liquid chromatography-mass spectrometry (LC-MS).

### Quantitative LC-MS and data analysis

SILAC-labeled cells of TbEMC3 and TbEMC8 RNAi experiments, mixed in equal ratios (+/-Tet; induced for 3 and 3.5 days, respectively; 3 replicates each with label-switch), were lyzed in 8 M urea/50 mM NH_4_HCO_3_. Proteins were reduced, alkylated and digested with trypsin in solution as described previously^49^. Peptide mixtures were analyzed by LC-MS using an Orbitrap Elite mass spectrometer (Thermo Fisher Scientific, Bremen, Germany) coupled to an UltiMate 3000 RSLCnano HPLC system (Thermo Fisher Scientific, Dreieich, Germany), which was operated with nanoEase™ M/Z Symmetry C18 precolumns (20 mm x 0.18 mm; flow rate, 10 µl/min; Waters) and a nanoEase™ M/Z HSS C18 T3 analytical column (250 mm x 75 µm; particle size, 1.8 µm; packing density, 100 Å; flowrate, 300 nl/min; Waters). Peptides were separated using a binary solvent system consisting of 0.1% formic acid (solvent A) and 30% acetonitrile/50% methanol/0.1% formic acid (solvent B). Following loading and concentration for 5 min at 7% B, peptides were eluted applying the following gradient: 7 - 52% B in 255 min, 52 - 65% B in 70 min, 65 - 95% B in 5 min and 5 min at 95% B.

The settings for the acquisition of mass spectrometric data in data-dependent mode were as follows: mass range, *m/z* 370 - 1,700; resolution, 120,000 (at *m/z* 400), target value, 1 × 10E6 ions; maximum injection time, 200 ms for MS survey scans; TOP25 method for low energy collision-induced dissociation of multiply charged peptides in the linear ion trap; normalized collision energy, 35%; activation q, 0.25; activation time, 10 ms; target value, 5,000 ions; maximum injection time, 150 ms; dynamic exclusion time, 45 s.

For protein identification and relative quantification, MaxQuant/Andromeda^60,61^ (version 1.6.5.0) was used. Mass spectrometric raw data were searched against a fasta file containing all entries for TREU927, downloaded from the TriTryp database (version 8.1; https://tritrypdb.org/). MaxQuant default settings were applied for data processing, except that proteins were identified and quantified based on ≥1 unique peptide and ≥1 ratio count, respectively. Carbamidomethylation of cysteine was set as fixed modification, acetylation of N-termini and oxidation of methionine were considered as variable modifications, and Lys8 and Arg10 were set as heavy labels. The options “requantify” and “match between runs” were enabled. The mean log_2_ of normalized protein abundance ratios (+/-Tet) was calculated and p-values (n = 3) were determined using a two-sided Student’s t-test. For information about proteins identified and quantified, see Table S9.

## Acknowledgments

The work was funded by Swiss National Science Foundation grants 169355 (to PB) and 175563 (to AS) and in part by the NCCR “RNA & Disease” funded by the Swiss National Science Foundation (to AS) and the Deutsche Forschungsgemeinschaft (DFG, German Research Foundation) project ID 403222702/SFB 1381 (to BW) and Germany’s Excellence Strategy CIBSS – EXC-2189 – project ID 390939984 (to BW). AI thanks S. Iyer and P.G. Balan for their support. BW thanks B. Knapp and W.D. Mühlhäuser for support in MS data acquisition and analysis. PB thanks R. Plant, M. Bütikofer and L. Apitzsch for their support.

## Notes

### Competing Interest Statement

The authors have declared no competing interest.

### Summary of Updates

This version of the manuscript now contains new mass spectrometry data and considers the reviewers' comments.

