## Supplemental Material 1 for "The endoplasmic reticulum membrane protein complex localizes to the mitochondrial - endoplasmic reticulum interface and its subunits modulate phospholipid biosynthesis in *Trypanosoma brucei*"

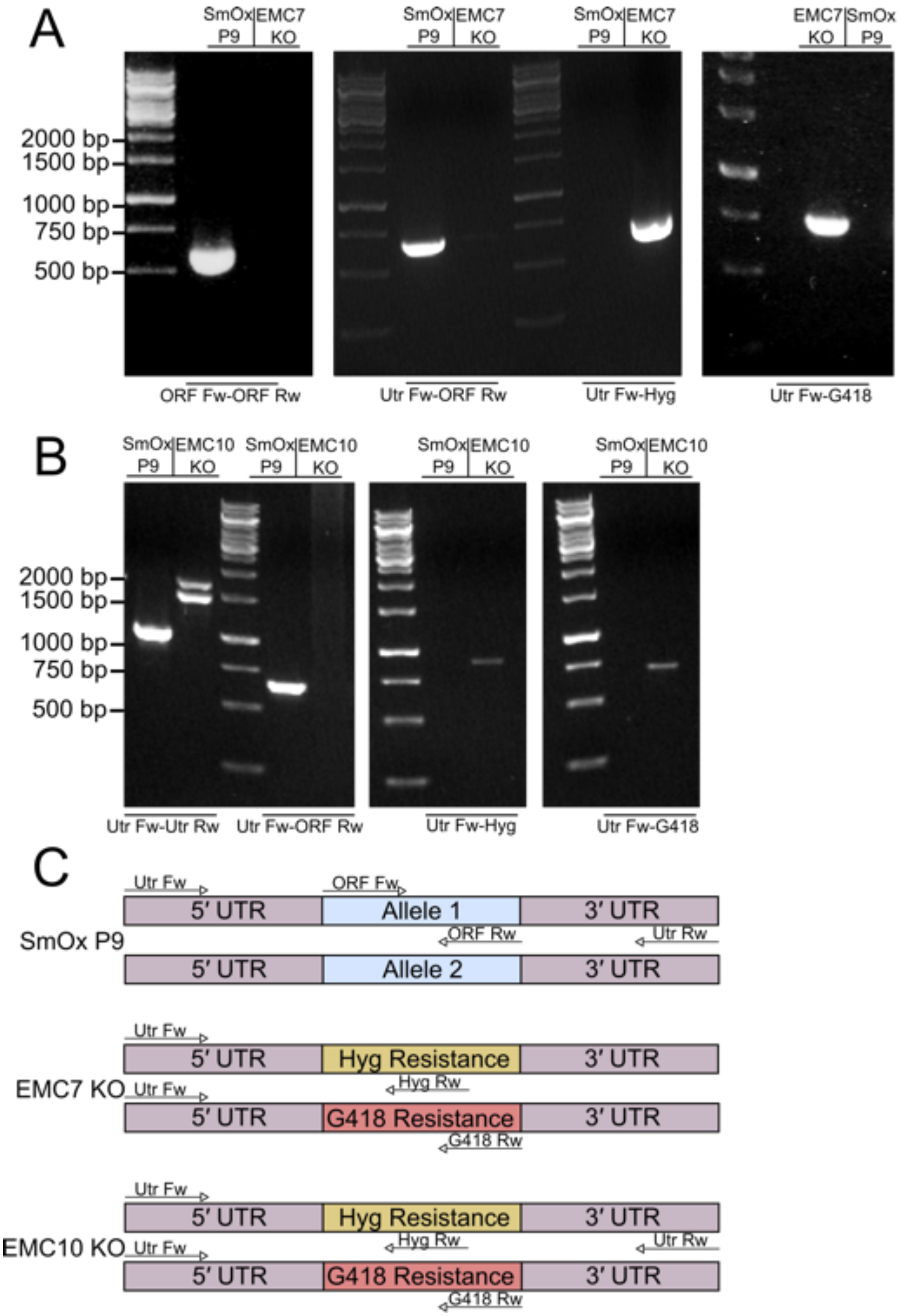


Figure S1. Verification of TbEMC-KO genotypes. gDNA extracted from parental (SmOx P9), TbEMC7-KO (A) and TbEMC10-KO (B) parasites was analyzed by PCR using the primer pairs indicated in C. The TbEMC alleles and hygromycin (Hyg) and geneticin (G418) resistance cassettes are indicated.

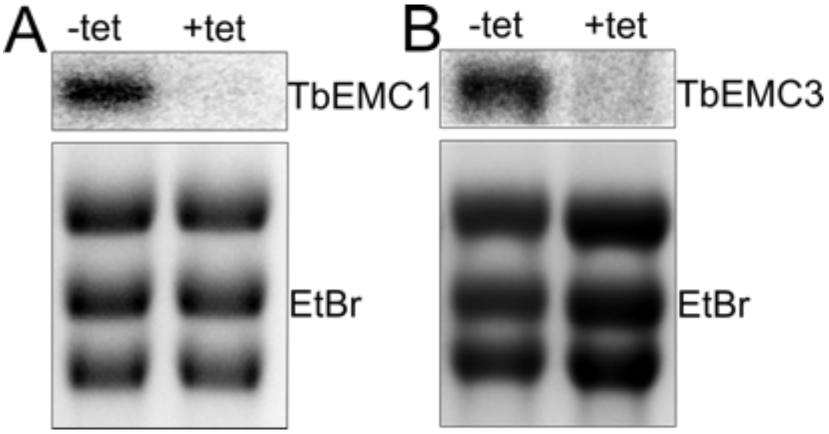


Figure S2. Verification of TbEMC mRNA levels. Total RNA extracted from parasites before (-tet) and after 3 days of induction of RNAi (+tet) was analyzed by Northern blotting using $^{32}P$-labeled probes against the ORFs of TbEMC1 (A) or TbEMC3 (B). Ethidium bromide-stained cytosolic rRNA served as loading control.

---

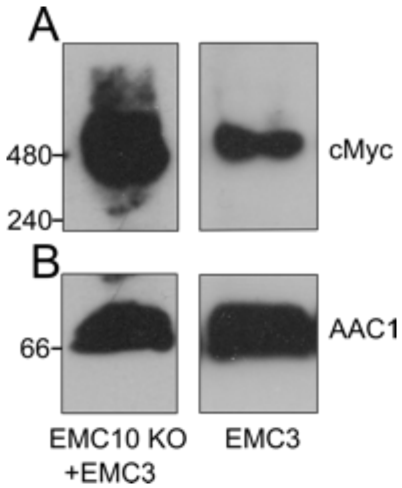


Figure S3. Analysis of TbEMC complex stability. Protein extracts from TbEMC10-KO (left lanes) and control (right lanes) parasites expressing *in situ* cMyc-tagged TbEMC3 were analyzed by native PAGE and immunoblotting using anti-cMyc (A) and anti-AAC1 (B, loading controls) antibodies. AAC1, ADP/ATP-carrier protein 1. Molecular mass markers (in kDa) are indicated in the left margin.

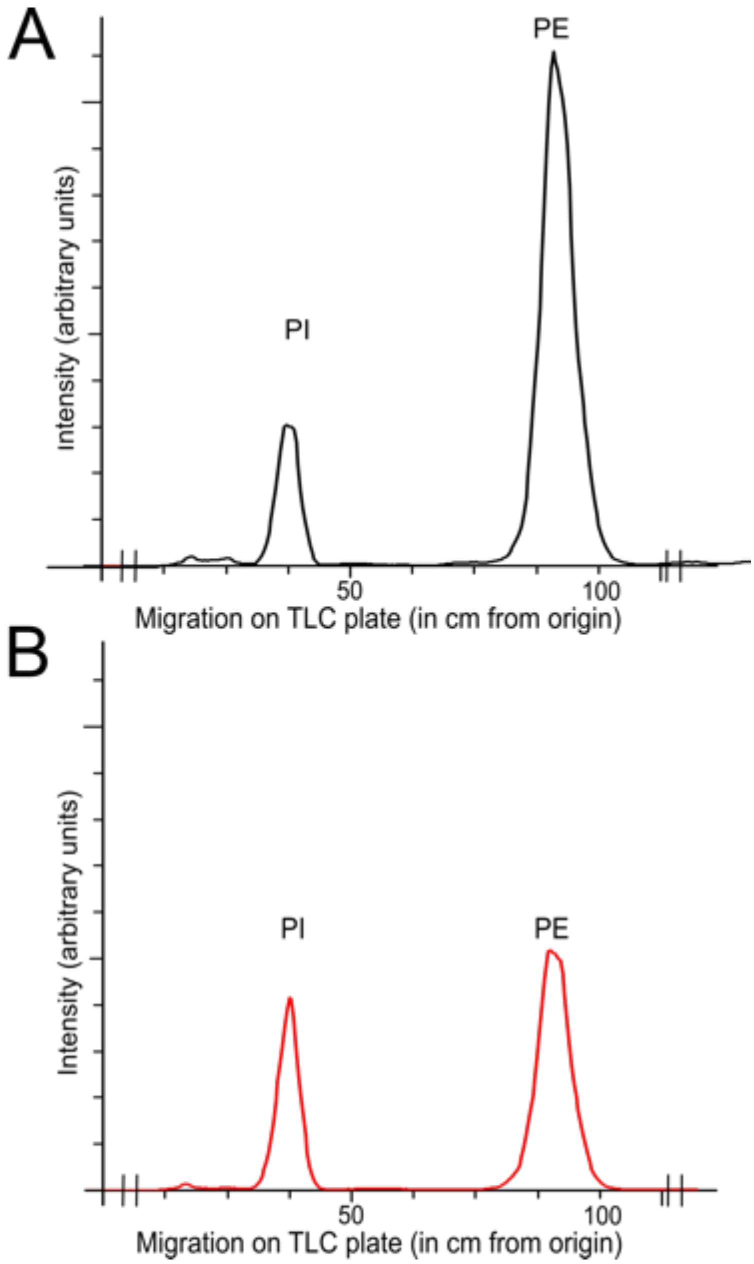


Figure S4. *De novo* synthesis of [$^3$H]-PE and [$^3$H]-PI in TbEMC8 RNAi cells. Parasites were cultured in the absence (A) or the presence (B) of tetracycline to maintain or ablate, respectively, TbEMC8 expression and incubated with [$^3$H]-ethanolamine and [$^3$H]-inositol (as control) for 4 h. Phospholipids were extracted and separated by TLC and incorporation of radioactivity into the different phospholipid classes was analyzed by radioisotope scanning.

Table S5. Primers used for generating *in-situ* tagged TbEMC cell lines.

| No. | Primer Name | Sequence |
|---|---|---|
| 1. | TbEMC1 fwd | ATGCGTAAAGGTGTGAGACAACTGTGGCGAGGGCCCCCCCTCG |
| 2. | TbEMC1 rev | CATCGCAACCAAGGCGACGACATGTTAATCGGCCGCTCTAGAACTAGTGG |
| 3. | TbEMC1 sgRNA | GAAATTAATACGACTCACTATAGGCCGAGAAGTCAATATCTCTCGTTTTAGAGCTAGAAATAGC |
| 4. | TbEMC2 fwd | GCACTGGAGGCGATGCTACATCGTGATGATGGGCCCCCCCTCG |
| 5. | TbEMC2 rev | CGCACACGCTTCTTTCACCCATCCCAACCCGGCCGCTCTAGAACTAGTGG |
| 6. | TbEMC2 sgRNA | GAAATTAATACGACTCACTATAGGCAGCTGCTTCTCAGCAGAACGTTTTAGAGCTAGAAATAGC |
| 7. | TbEMC3 fwd | GATGCCCCCCGTTTACTCCTTGAAGGTAAAGGGCCCCCCCTCG |
| 8. | TbEMC3 rev | CCTTTCCTGAAATTTGGCTTCAAACAGCCA GGCCGCTCTAGAACTAGTGG |
| 9. | TbEMC3 sgRNA | GAAATTAATACGACTCACTATAGGGCAGCAGCAGTGTACTTCAAGTTTTAGAGCTAGAAATAGC |
| 10. | TbEMC4 fwd | AAGGAGCAGAGCATGGGCACATTTTTAGTGGGCCCCCCCTCG |
| 11. | TbEMC4 rev | CTTAACCACCACAAACTAGCGCCAAAACCGGGCCGCTCTAGAACTAGTGG |
| 12. | TbEMC4 sgRNA | GAAATTAATACGACTCACTATAGGTTGTATACATGTGTGTGTATGTTTTAGAGCTAGAAATAGC |
| 13. | TbEMC5 fwd | ATTCGAGCATCAACAGGGAAGAAACATGCGGGGCCCCCCCTCG |
| 14. | TbEMC5 rev | GATCCAAAACTGCAACAGTCTCTGTTGCCGGGCCGCTCTAGAACTAGTGG |
| 15. | TbEMC5 sgRNA | GAAATTAATACGACTCACTATAGGTTATTTGTTTGTGTAGTCGGGTTTTAGAGCTAGAAATAGC |
| 16. | TbEMC6 Fwd | CGCGGTGATAGCACGGGTGTAGAAAATAGTGTATAATGCAGACCTGCTGC |
| 17. | TbEMC6 Rev | AAAAGTGAGCATGCAACCACGTCCAAACATACTACCCGATCCTGATCCAG |
| 18. | TbEMC6 SgRNA | GAAATTAATACGACTCACTATAGGATTTTTTTTTCTGTTACGAGTTTTAGAGCTAGAAATAGC |
| 19. | TbEMC7 Fwd | ACTGCGGCGGGACACGAGAGGAAGCACAAGGGGCCCCCCCTCG |
| 20. | TbEMC7 Rev | CCCTTACCCACTACCCAACCACAAATTCCAGGCCGCTCTAGAACTAGTGG |
| 21. | TbEMC7 SgRNA | GAAATTAATACGACTCACTATAGGGTTGTTGGTAGTTATGATAGGTTTTAGAGCTAGAAATAGC |
| 22. | TbEMC8 Fwd | TTGAACTACCTCGACCAAACAGTCTCCGCCGGTACCGGGCCCCCC |
| 23. | TbEMC8 Rev | TGAGGTCAAAGAAAAGAACGAAAGAAGGAATGGCGGCCGCTCTAGAAC |
| 24. | TbEMC8 SgRNA | GAAATTAATACGACTCACTATAGGAGGAAAGGAAACTAAAGAAGGTTTTAGAGCTAGAAATAGC |
| 25. | TbEMC10 Fwd | TTACTTCAGGGCATTTTCGCGCCTAAGCAGGGGCCCCCCCTCG |
| 26. | TbEMC10 Rev | CACCACACAAATGCACACACATAATCCCCTGGCCGCTCTAGAACTAGTGG |
| 27. | TbEMC10 SgRNA | GAAATTAATACGACTCACTATAGGCAGGAGGGCGCGGGGGAGAGGTTTTAGAGCTAGAAATAGC |

Table S6. Primers used for generating TbEMC KO and TbEMC RNAi cell lines.

| **No.** | **Primer Name** | **Sequence** |
|---|---|---|
| 1. | TbEMC7 KO Fwd | CGCGGTGATAGCACGGGTGTAGAAAATAGTGTATAATGCAGACCTGCTG |
| 2. | TbEMC7 KO Rev | CCCTTACCCACTACCCAACCACAAATTCCACCGGAACCACTACCAGAACC |
| 3. | TbEMC7 KO 5`SgRNA | GAAATTAATACGACTCACTATAGGATTTTTTTTTCTGTTACGAGTTTTAGAGCTAGAAATAGC |
| 4. | TbEMC7 KO 3`SgRNA | GAAATTAATACGACTCACTATAGGGTTGTTGGTAGTTATGATAGGTTTTAGAGCTAGAAATAGC |
| 5. | TbEMC10 KO Fwd | AAAGTCGGGTAAGTCAGGTACCTCATTTTCGTATAATGCAGACCTGCTGC |
| 6. | TbEMC10 KO Rev | CACCACACAAATGCACACACATAATCCCCTCCGGAACCACTACCAGAACC |
| 7. | TbEMC10 KO 5`SgRNA | GAAATTAATACGACTCACTATAGGCCTTTCACAATAAAGTGTGAGTTTTAGAGCTAGAAATAGC |
| 8. | TbEMC10 KO 3` SgRNA | GAAATTAATACGACTCACTATAGGCAGGAGGGCGCGGGGGAGAGGTTTTAGAGCTAGAAATAGC |
| 9. | TbEMC1 RNAi Fwd | CCCAAGCTTGGGGGATCCGTTTGAAACCGAGGTTGGCC |
| 10. | TbEMC1 RNAi Rev | CTAGTCTAGACTCGAGAGAGGTGCGCACGTAAAAGA |
| 11. | TbEMC3 RNAi Fwd | ATCTGAAGCTTGGATCCTGTACTTCAATGGCTGTTTG |
| 12. | TbEMC3 RNAi Rev | GCAGATCTAGACTCGAGACTACCCTATCCCTCAGCTC |
| 13. | TbEMC8 RNAi Fwd | CCCAAGCTTGGATCCCACAGCAGAAAATCCGGTG |
| 14. | TbEMC8 RNAi Rev | CCCTCTAGACTCGAGGGTCGAGGTAGTTCAATTGTAC |

Table S7. Putative TbEMC3 interaction partners.

| Gene ID | Annotation | Fold-Change (log2) | adjusted p-Value |
|---|---|---|---|
| Tb927.10.4760 | EMC3 | 6.3 | 4.6E-06 |
| Tb927.11.14130 | Ribosomal protein L18, putative | 5.7 | 6.4E-06 |
| Tb927.8.1570 | EMC8 | 5.6 | 1.5E-05 |
| Tb927.11.9730 | 60S ribosomal protein L34, putative | 5.4 | 2.8E-05 |
| Tb927.7.6260 | EMC2 | 5.3 | 6.6E-06 |
| Tb927.9.15420 | 60S ribosomal protein L32 | 5.1 | 1.2E-05 |
| Tb927.9.3990 | Ribosomal protein S7, putative | 5.1 | 1.5E-05 |
| Tb927.4.2180 | 60S ribosomal protein L35a, putative | 5.1 | 5.9E-05 |
| Tb927.10.5620 | Fructose biphosphate aldolase, glycosomal | 4.4 | 4.9E-05 |
| Tb927.10.190 | 40S ribosomal protein S6, putative | 4.4 | 1.5E-05 |
| Tb927.9.5860 | Tryparedoxin peroxidase | 4.0 | 6.4E-06 |
| Tb927.10.3280 | 60S ribosomal protein L38, putative | 4.0 | 1.5E-05 |
| Tb927.10.5370 | 40S ribosomal protein S10, putative | 3.9 | 1.5E-05 |
| Tb927.9.6070 | 40S ribosomal protein S3, putative | 3.8 | 9.0E-06 |
| Tb927.10.10590 | Histone H2B, putative | 3.8 | 4.9E-05 |
| Tb927.3.1120 | GTP-binding nuclear protein rtb2, putative | 3.6 | 1.5E-05 |
| Tb927.9.12240 | 60S ribosomal subunit protein, L31, putative | 3.6 | 2.1E-04 |
| Tb927.2.6090 | 60S ribosomal protein L44 | 3.5 | 8.2E-05 |
| Tb927.11.15900 | 60S ribosomal protein L27, putative | 3.5 | 1.2E-05 |
| Tb927.10.12840 | Mitochondrial carrier protein 12 | 3.4 | 2.1E-04 |
| Tb927.7.5180 | 60S ribosomal protein L23a, putative | 3.3 | 1.9E-04 |
| Tb927.11.4820 | 60S ribosomal protein L17, putative | 3.2 | 3.8E-04 |
| Tb927.9.15210 | Ribosomal protein L36, putative | 3.0 | 1.5E-05 |
| Tb927.11.10910 | 40S ribosomal protein SA, putative | 2.7 | 2.6E-03 |
| Tb927.10.5480 | 60S ribosomal protein L24, putative | 2.7 | 4.9E-05 |
| Tb927.7.1750 | 60S ribosomal protein L7, putative | 2.6 | 4.9E-05 |
| Tb927.11.8200 | Ribosomal protein S26, putative | 2.5 | 3.7E-05 |
| Tb927.11.9710 | 60S ribosomal protein L10a, putative | 2.4 | 3.8E-04 |
| Tb927.10.1100 | 60S ribosomal protein L9, putative | 2.4 | 2.5E-05 |
| Tb927.11.14020 | Nuclear RNA binding domain 2 | 2.3 | 1.9E-04 |
| Tb927.5.1110 | 60S ribosomal protein L8, putative | 2.2 | 1.1E-03 |
| Tb927.9.11410 | 60S ribosomal protein L23, putative | 2.2 | 1.4E-04 |
| Tb927.2.5910 | 40S ribosomal protein S13, putative | 2.1 | 1.6E-04 |
| Tb927.7.1050 | 40S ribosomal protein S16, putative | 2.1 | 8.2E-05 |

*TbEMC subunits are indicated in yellow and ribosomal proteins in blue.*

Table S8. Putative TbEMC5 interaction partners.

| Gene ID | Annotation | Fold-Change (log2) | adjusted p-Value |
|---|---|---|---|
| Tb927.8.1570 | EMC8 | 8.7 | 7.8E-08 |
| Tb927.7.6260 | EMC2 | 8.3 | 2.1E-07 |
| Tb927.10.4760 | EMC3 | 8.0 | 2.1E-07 |
| Tb927.4.2320 | EMC5 | 5.1 | 1.8E-05 |
| Tb927.3.1120 | GTP-binding nuclear protein rtb2, putative | 3.7 | 2.6E-04 |
| Tb927.11.11680 | 2-oxoglutarate dehydrogenase E2 component, put. | 3.4 | 2.6E-04 |
| Tb927.9.11600 | Glycosomal membrane protein | 3.0 | 5.0E-05 |
| Tb927.4.590 | EMC1 | 2.8 | 3.8E-03 |
| Tb927.6.2600 | EMC4 | 1.9 | 8.4E-04 |

*TbEMC subunits are indicated in yellow.*
